## Supplemental_Figures for "A distinct assembly pathway of the human 39S late pre-mitoribosome"

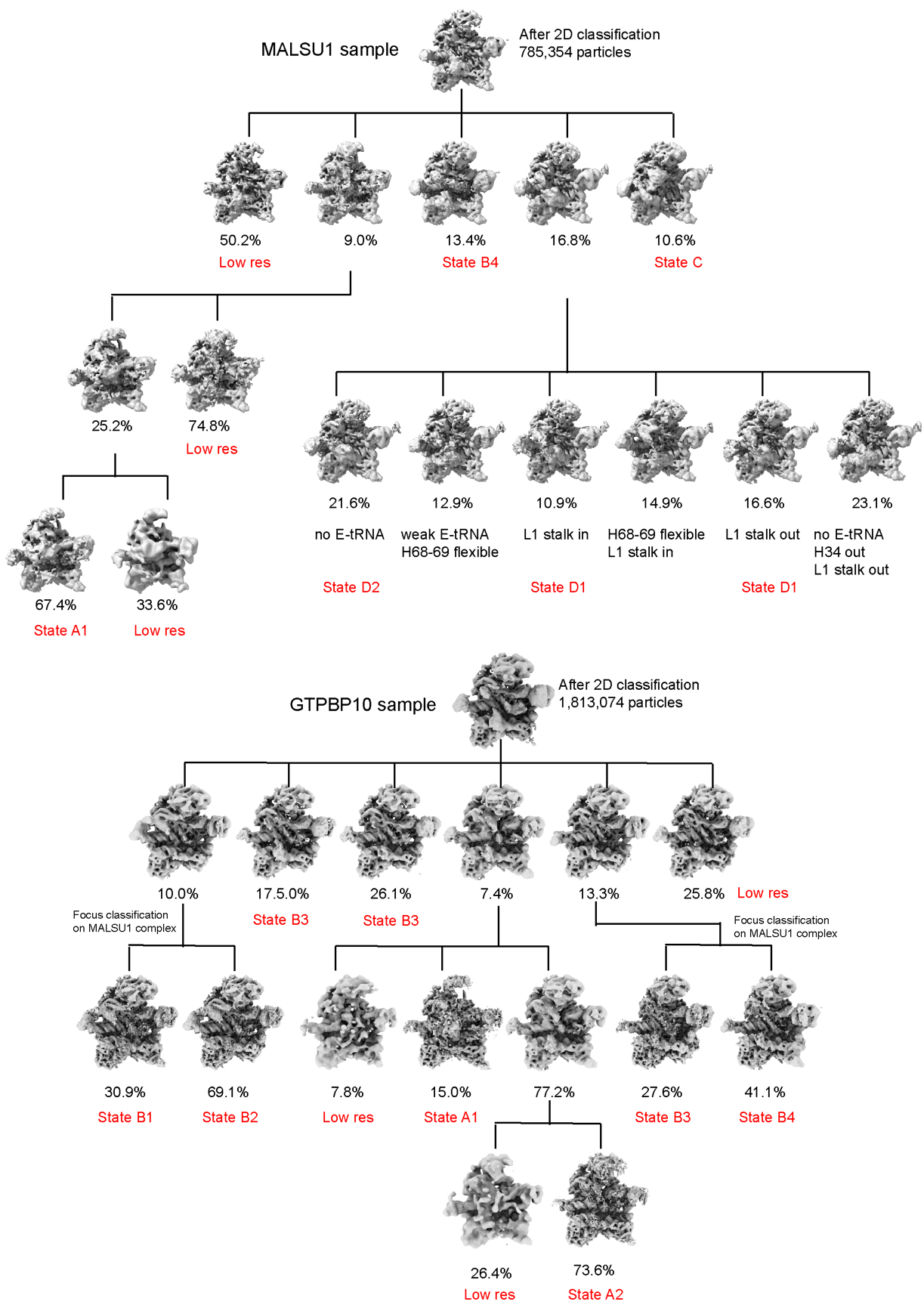

**Supplementary Figure 1 | Cryo-EM analysis of the MALSU1 and GTPBP10 samples**

Schematic breakdown of the cryo-EM analysis for the datasets collected from the MALSU1 and GTPBP10 samples. Low res: low resolution classes.

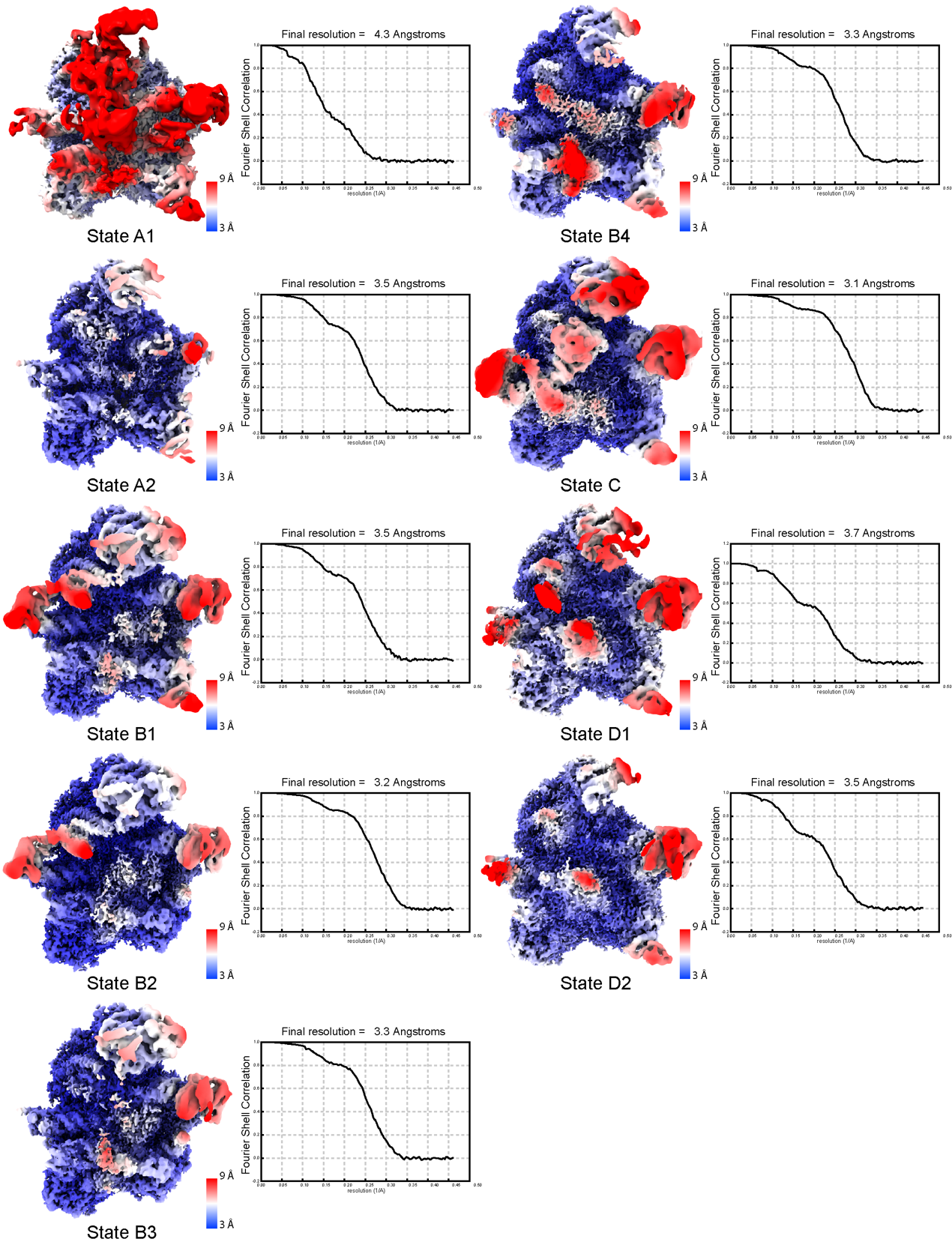

**Supplementary Figure 2 | Local resolution distribution and Fourier shell correlation plot of different states.**

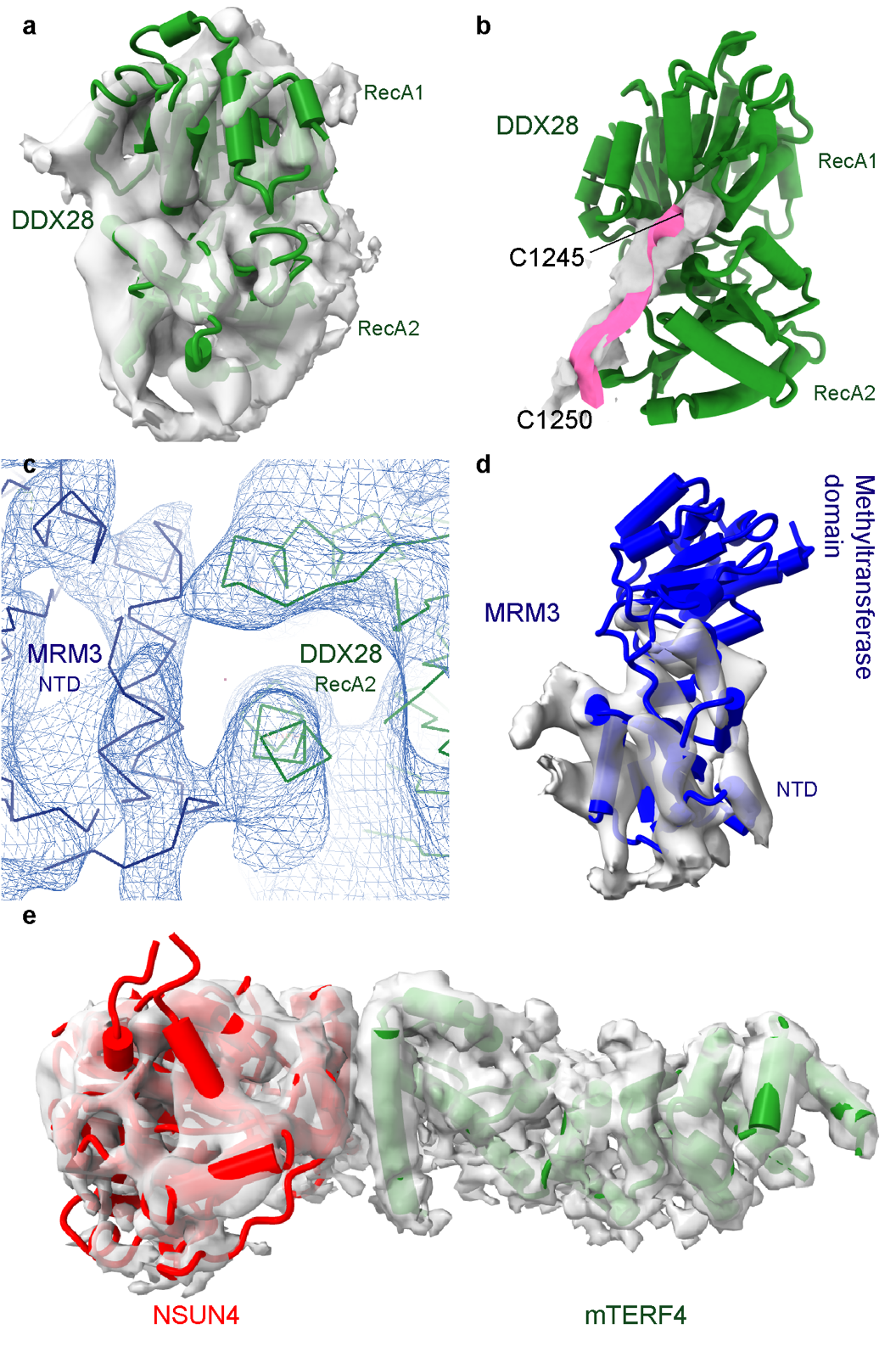

**Supplementary Figure 3 | Structure of DDX28, MRM3, NSUN4 and mTERF4**

**a,** DDX28 from state A1 is surrounded by gray density to show the rigid body fit. **b,** A close up view of the RNA binding tunnel of DDX28 (between RecA1 and RecA2). There is a clear RNA density inside the tunnel corresponding to 16S rRNA region (1245-1250). **c,** The direct interaction between the NTD of MRM3 and the RecA2 domain of DDX28. The map corresponding to these regions is shown as a mesh. **d,** MRM3 from state A1 is surrounded by gray density to show the rigid body fit of the N terminal domain (NTD). **e,** The state C NSUN4-mTERF4 complex (red and green model respectively) is shown with gray density.

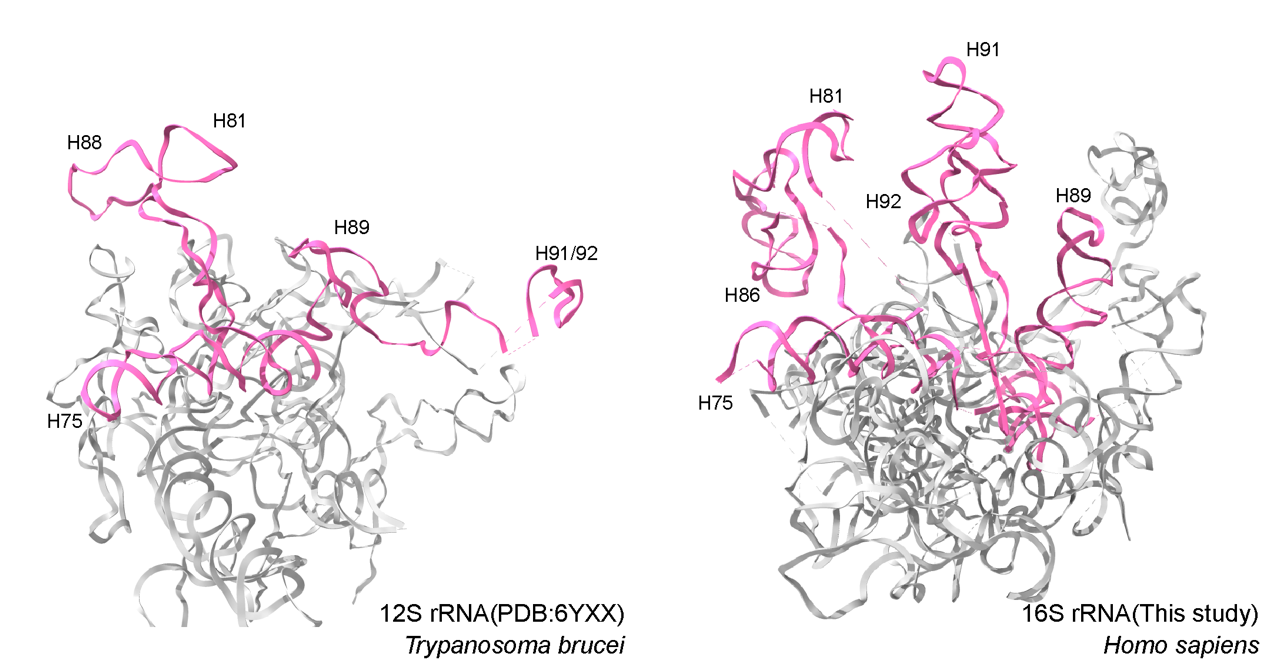

**Supplementary Figure 4 | Comparison of the immature rRNA**

Interspecies differences in folding of domain V of the human 16S rRNA and homologous counterpart of the 12S rRNA from *Trypanosoma brucei.* The immature domain V from both species are colored in pink, the residual 12S and 16S rRNA domains are colored in gray.
